# TGFam-Finder: An optimal solution for target-gene family annotation in eukaryotic genomes

**DOI:** 10.1101/372433

**Authors:** Seungill Kim, Kyeongchae Cheong, Jieun Park, Myung-Shin Kim, Ji-Hyun Kim, Min-Ki Seo, Sun-Ho Kwon, Yong-Min Kim, Namjin Koo, Kwang-Soo Kim, Nuri Oh, Ki-Tae Kim, Jongbum Jeon, Hyunbin Kim, Yoon-Young Lee, Kee Hoon Sohn, Honour C McCann, Sang-Kyu Ye, Kyung-Soon Park, Yong-Hwan Lee, Doil Choi

## Abstract

Whole genome annotation errors that omit essential protein-coding genes hinder further research. We developed Target Gene Family Finder (TGFam-Finder), an optimal tool for structural annotation of protein-coding genes containing target domain(s) of interest in eukaryotic genomes. Large-scale re-annotation of 100 publicly available eukaryotic genomes led to the discovery of essential genes that were missed in previous annotations. An average of 117 (346%) and 148 (45%) additional FAR1 and NLR genes were newly identified in 50 plant genomes. Furthermore, 117 (47%) additional C2H2 zinc finger genes were detected in 50 animal genomes including human and mouse. Accuracy of the newly annotated genes was validated by RT-PCR and cDNA sequencing in human, mouse and rice. In the human genome, 26 newly annotated genes were identical with known functional genes. TGFam-Finder along with the new gene models provide an optimized platform for unbiased functional and comparative genomics and comprehensive evolutionary study in eukaryotes.

## Introduction

The recent emergence of long read sequencing approaches such as single molecule real-time sequencing and nanopore technology enables near-perfect assemblies of even enormous genomes like that of the salamander^1^ (genome size = 32 Gb, contig N50 = 300 kb) and the improvement of existing assemblies for complex genomes^2-4^. Structural gene annotation of protein-coding sequences is a post-assembly process that is essential for further research^5^. To date, hundreds of plant and animal genome resources have been released in public databases and used for subsequent functional genomics analyses, evolutionary analyses, and biotechnology applications. However, there are continuous reports of annotation errors, including imperfect gene models and missing functional genes^6-8^. These reports demonstrate that the accidental omission of essential genes that are correctly located in assembled genomes but not annotated in the gene model can ultimately generate biases in downstream studies.

Pre-existing gene models are continuously being improved using manual, computational, and experimental analyses for model species such as human, mouse, and Arabidopsis. Since the Gene Encyclopedia of DNA Elements (GENCODE) project was initiated, a total of 25 and 17 updates for whole gene models of human and mouse have been accomplished, respectively (see URLs). In plant, The Arabidopsis Information Resource (TAIR) is maintaining genomic resources for the model plant *A. thaliana*, currently providing the 11^th^ updated gene model, (Araport 11, see URLs). In addition, groups conducting genome sequencing projects of major animal and plant species continuously improve the quality of their gene models (see URLs). However, the majority of published gene models remain inaccurate and likely incomplete, and have not been updated beyond the initial version. In general, the potential function of a gene is predicted based on the identification of conserved domains or motifs. Studies focusing on specific genes or families often begin working with annotated gene models by identifying those genes of interest that contain the appropriate target domain(s) or motifs^8,9^. Performing new annotation and improving existing gene models require huge inputs of human labor and computational resources. Therefore, researchers have designed novel approaches to identify specific genes or gene families^7, 9-11^. For example, Jupe et al. developed resistance gene enrichment and sequencing (RenSeq), a high-throughput sequencing method for the selective capture and sequencing of nucleotide-binding and leucine-rich-repeat (NLR) genes without whole genome sequencing^9^. Although these methods enable detection of candidate regions containing target genes, further annotation to determine accurate gene structure in candidate regions remains a bottleneck.

Here, we present Target Gene Family Finder (TGFam-Finder), an optimal tool that allows automatic annotation of all protein-coding genes containing specific target domain(s) in assembled genomes. We evaluated TGFam-Finder through the massive re-annotation of far-red-impaired response 1 (FAR1) transcription factor and nucleotide-binding and leucine-rich-repeat (NLR) gene families in 50 plant genomes, as well as Cys2-His2 zinc finger (C2H2 zinc finger) and homeobox transcription factor gene families in 50 animal genomes. A large number of missing target genes in the pre-existing gene models were newly identified during re-annotation with TGFam-Finder. In particular, 26 newly annotated genes that were omitted in the existing human gene model had identical sequences with known functional genes. We validated the accuracy and expression of these newly annotated genes in human, mouse, and rice by performing RT-PCR and cDNA sequencing analyses. Our analyses demonstrate the efficiency of TGFam-Finder for users, including bench-based researchers given notably reduced annotation time using a desktop computer. TGFam-Finder, a domain search-based gene annotation tool could provide an optimal solutions for target-gene family annotation in functional, comparative, and evolutionary studies.

## Results

### Conceptual overview of TGFam-Finder

We designed TGFam-Finder as an unbiased annotation tool to identify any target-gene family of interest in assembled genomes. TGFam-Finder was developed for an audience including novice bioinformaticians, incorporating ease of use from installation to completion of structural annotation. To this, we provide additional tool packages enabling the automatic installation of prerequisite tools for further structural gene annotation using TGFam-Finder without any manual configuration (Supplementary Fig. 1 and Online Methods)

An automatic annotation process using TGFam-Finder consists of the following three steps : (1) genome-wide identification of target regions containing specific target-gene sequences of interest, (2) structural annotation of the target regions using available proteins, transcriptomes and *ab-initio* prediction, and (3) construction of the final gene model (Fig. 1). One of the distinct features of TGFam-Finder is the extraction of target regions containing sequences of target-gene family. To reduce annotation time and unnecessary computation, TGFam-Finder identifies all genomic regions containing domain(s) of the target genes using HMMER^12^ from six-frame translated genome sequences. The target regions are determined after masking unnecessary sequences as ‘*X*’, except for the identified genomic regions and their flanking sequences (Fig. 1). Then, structural annotation of target regions is performed to generate the initial gene model through serial processes of protein mapping, transcriptome annotation, and *ab-initio* prediction (Fig. 1). On the basis of the evidence gathered in the previous steps, TGFam-Finder combines the initial gene models and determines the final gene model of target families (Fig.1).

**Figure 1.**
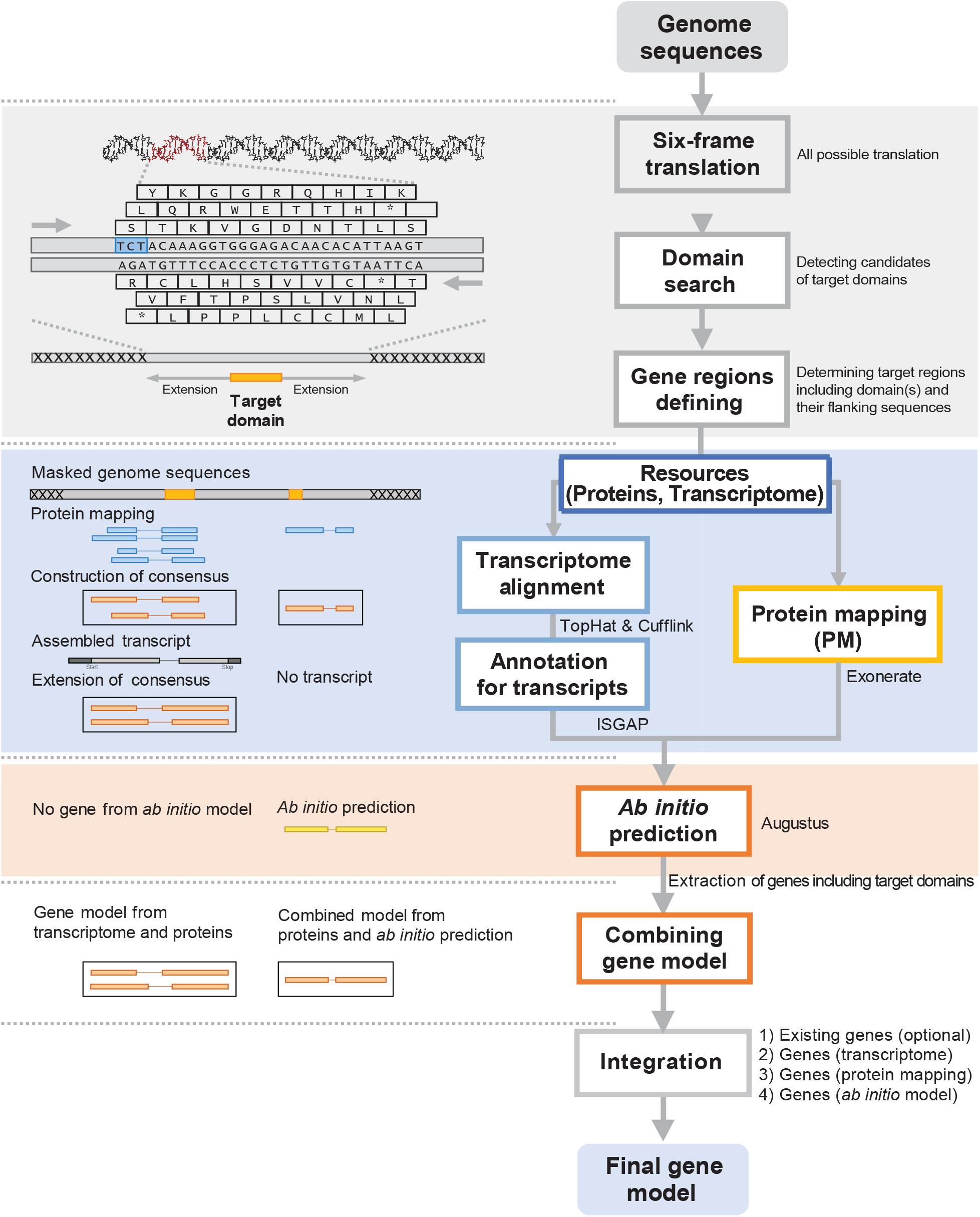
Annotation process of TGFam-Finder. An automatic process for new identification of target-gene families using TGFam-Finder is depicted. The diagram shows serial processes starting from six-frame translation to integration of the gene model. The gray block of the diagram describes determination of target regions containing target domain(s) and their flanking sequences for further annotation. The blue and pink blocks indicate structural annotation using proteins and transcriptomes, and the *ab-initio* method, respectively. Names of representative tools^21-24^ for structural annotation are given in the blue and pink blocks. Initial gene models are integrated from the structural annotation as depicted in the white block.

TGFam-Finder improves upon pre-existing gene models through identification of missing essential genes, providing an unbiased model of target-gene families. To evaluate TGFam-Finder, we collected genomic data for 50 plants and 50 animals, including assemblies, annotated genes, and transcriptome data from public databases (Supplementary Table 1-2). In the plant genomes, we searched for the FAR1 transcription factor family that modulates phytochrome A signaling^13^ and NLR gene family that typically contains plant cytoplasmic immune receptor genes^14^ as target-gene families. In the animal genomes, we searched for the C2H2 zinc finger proteins, the largest transcription factor family, which is involved in functions such as sequence-specific DNA-binding and protein-protein interaction^15^, and the homeobox transcription factor family which primarily induces cellular differentiation by transcriptionally controlling co-regulated gene expression cascades^16^.

### FAR1 and NLR annotation in plant genomes

We re-annotated FAR1 and NLR genes in 50 plant genomes using TGFam-Finder (Fig. 2, Supplementary Fig. 2-3 and Online Methods). Only 1.3% and 1.9% of the plant genomes (average genome length 1,127 Mb) were determined as target regions for the re-annotation of FAR1 and NLR, respectively (Supplementary Fig. 2 and Supplementary Table 3). On average, 34 FAR1 and 327 NLR genes considering representative loci were identified in previously annotated gene models of the 50 plant genomes (Supplementary Table 3). In addition to these, we identified 117 (346%) and 148 (45%) new FAR1 and NLR genes using TGFam-Finder, respectively, indicating that only 23% (34 of 151) of FAR1s and 69% (327 of 475) of NLRs were annotated in previous studies (Fig. 2a, Supplementary Fig. 3a, and Supplementary Table 3). Specifically, 104 (89%) and 108 genes (73%) of the newly annotated FAR1 and NLR genes were located in genomic regions without any genes in the existing models (Supplementary Fig. 4 and Supplementary Table 4). In addition, TGFam-Finder found intact gene structures in regions containing previously annotated partial FAR1 and NLR genes that have no start or stop codons (Supplementary Table 3). TGFam-Finder newly annotated 31 FAR1s (26%) and 69 NLRs (43%) based on protein or transcriptome evidences in the new gene models (Supplementary Table 5).

**Figure 2.**
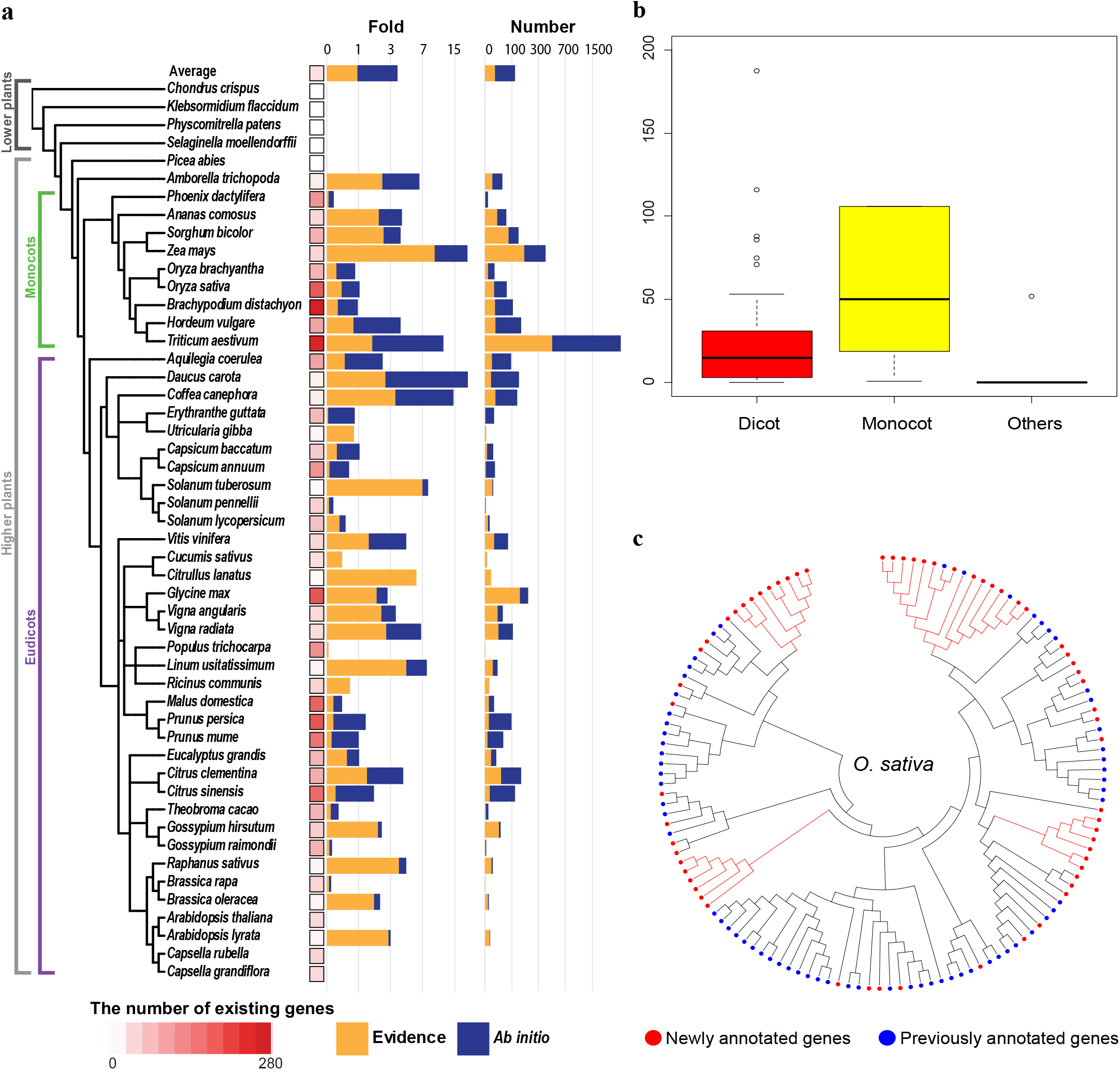
Re-annotation of FAR1 genes in 50 plant genomes. (a) Heat map indicates the number of existing FAR1 genes in the representative loci of 50 plant genomes. Bar graphs show the fold-increase of newly annotated genes compared with the number of existing genes (left) and the number of newly annotated genes (right). Colors in the bar represent the number of newly annotated genes from protein or transcriptome evidence (orange) and *ab-initio* model (navy blue). (b) Boxplot shows the distribution of the number of newly annotated genes in clusters without any existing genes. (c) Phylogenetic tree of newly annotated genes in the rice genome. Red and blue circles indicate newly annotated and previously annotated genes, respectively. Red branches indicate sublineages with multiple newly annotated genes including no copies or only a small number of previously annotated genes.

Compared to the number of genes in the existing gene models, we found a large number of new FAR1s in 18 of the 50 genomes, including 12 dicot and 5 monocot plants, which was more than a 3-fold increase, or more than 100 new FAR1s (Fig.2a). Specifically, 7 and 19 FAR1s were annotated in the existing model of carrot and maize genomes, but TGFam-Finder identified 142 and 383 new FAR1s in carrot and maize, respectively, a more than 20-fold increase relative to the number of existing genes (Fig. 2a and Supplementary Table 3). In the wheat genome, we detected more than 3,000 new FAR1 genes. For the new gene model of NLRs, we also found a number of genes in 14 plant genomes (9 dicot species), which is a more than 1-fold increase or more than 200 genes (Supplementary Fig. 3a and Supplementary Table 3). Although only three NLRs were annotated in existing gene model of *Selaginella* genome, 56 (>18-fold increase) more NLRs were correctly detected by TGFam-Finder in the same genome sequences (Supplementary Fig. 3a and Supplementary Table 3). Furthermore, we identified over 1,000 more NLRs in genome sequences of *Eucalyptus* and wheat, respectively. These results indicate that gene models for FAR1 and NLR were greatly improved using TGFam-Finder compared with the extremely biased existing models omitting many of those genes.

To study phylogenetic relationships of FAR1 and NLR in the 50 plant genomes, we performed gene clustering and phylogenetic analyses of the new gene models containing the newly annotated genes and the existing gene models (Fig. 2b, Supplementary Fig. 3b and Supplementary Table 6). The gene clustering analyses among the new gene models of 50 plant genomes revealed that an average of 48 (41%) new FAR1 genes and 20 (13%) new NLR genes did not cluster with existing genes (Supplementary Table 6). Overall, the number of FAR1s in new clusters was significantly larger in monocots than in dicots (Fig. 2b). These results indicate that a large number of the newly annotated genes, especially FAR1 in monocots, belonged to new subfamilies distinguished from the other subfamilies of the existing gene models. Furthermore, phylogenetic analyses of FAR1 and NLR in various plant genomes revealed that a large number of newly annotated genes were distinctly grouped in specific lineages. This means that certain gene clades were never annotated in earlier studies (Fig. 2c, Supplementary Fig. 3c and 4).

### Annotation of C2H2 zinc finger and homeobox genes in animal genomes

We performed re-annotation of C2H2 zinc finger and homeobox gene families in 50 animal genomes (Fig. 3 and Supplementary Fig. 5-6). A total of only 0.6% and 1.7% of the animal genomes (average genome size 1,652Mb) were used as target regions for the re-annotation of C2H2 zinc finger and homeobox genes, respectively (Supplementary Fig. 5 and Supplementary Table 7). An average of 117 (47%) and 22 (14%) additional C2H2 zinc finger and homeobox genes, respectively, were newly annotated in the 50 animal genomes (Supplementary Table 7). Specifically, 75% (88 and 17) of the new C2H2 zinc finger and homeobox genes did not overlap with any previously annotated genes, indicating that majority of the newly annotated genes were located in new chromosomal regions where no genes had been identified in earlier studies (Supplementary Table 8). We observed many partial C2H2 zinc finger and homeobox genes in existing gene models of several invertebrate genomes in contrast to the result of re-annotation of plant genomes. In that regions containing those partial genes, TGFam-Finder identified new or their intact gene structures (Supplementary Table 7).

**Figure 3.**
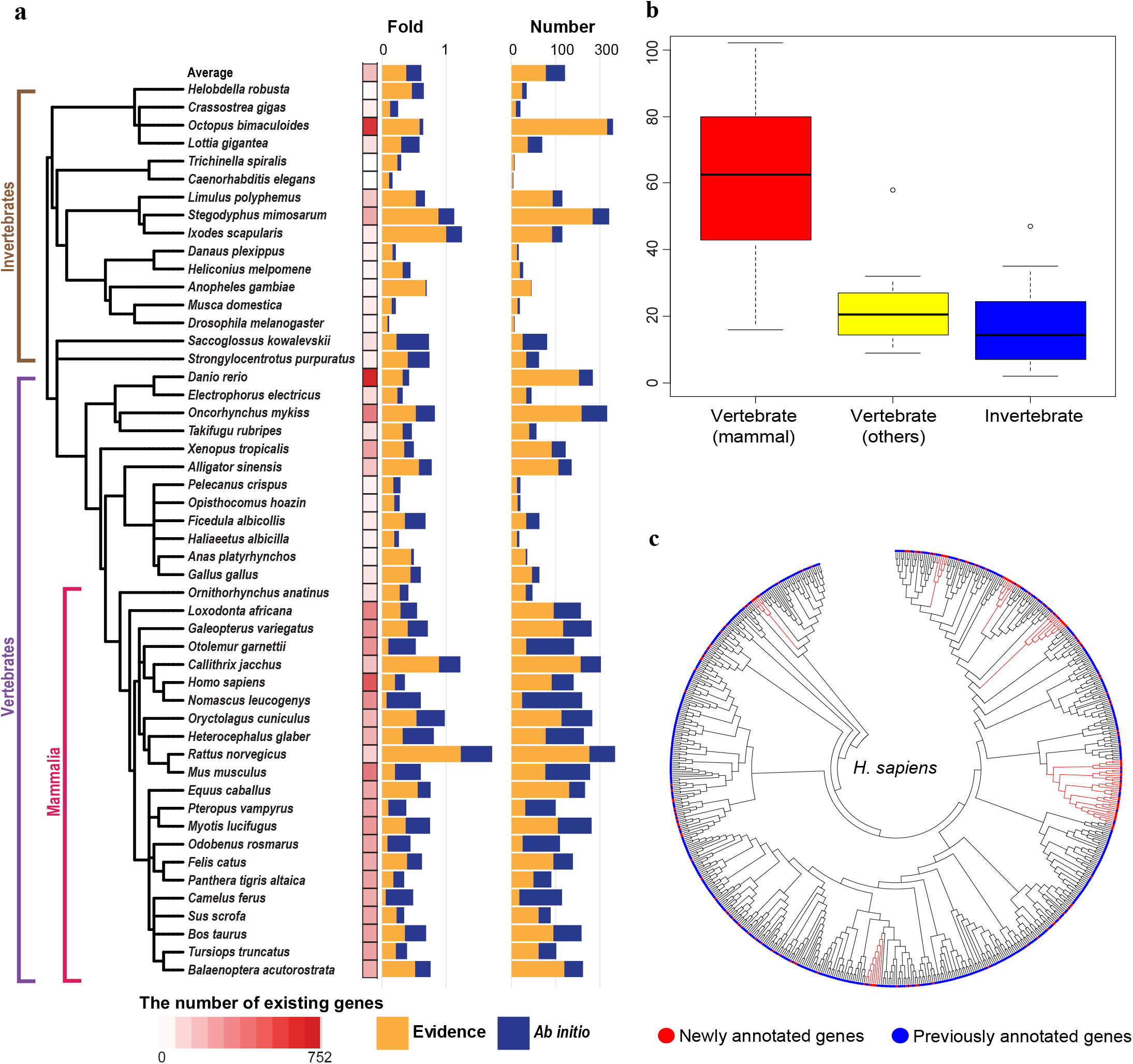
Newly annotated C2H2 zinc finger genes in 50 animal genomes. (a) Heat map and bar graphs represent the number of previously annotated genes in representative loci and the fold-increase and number of newly annotated genes, respectively. Orange bars indicate the number of genes generated from protein or transcriptome evidence. Navy blue bars represent *ab-initio* prediction. (b) The number of newly annotated genes in clusters without any previously identified genes is depicted. (c) Phylogenetic tree of new gene model in the human genome shows the phylogenetic relationship of those genes. The newly annotated and previously annotated genes are marked as red and blue circles, respectively.

Our analyses revealed that over half of the new C2H2 zinc finger (54%) and homeobox genes (66%) were annotated based on protein or transcriptome evidence, indicating that a great number of new genes were identified by obvious evidence (Supplementary Table 9). We found a number of new C2H2 zinc finger (> 1-fold increase compared to the number of existing genes or 100 additional copies) in 24 of the 50 animal genomes including 17 genomes of mammalian species (Supplementary Table 7-8). In particular, we observed 133 (23%) and 227 (47%) newly identified C2H2 zinc finger genes in human and mouse genomes, respectively, and 77 (33%) new homeobox genes in the human genome (Supplementary Table 7-8). Together with the re-annotation of FAR1 and NLR in plant genomes, these results confirm that TGFam-Finder significantly improved the existing gene models of C2H2 zinc finger and homeobox genes in the animal genomes by identifying new gene structures absent in the existing gene models.

Phylogenetic comparisons of C2H2 zinc finger and homeobox gene families using the new gene models revealed that the many (36%) newly annotated C2H2 zinc finger genes were clustered as unique clades, suggesting that those were derived from newly identified lineages that were absent in existing models (Fig. 3b, Supplementary Fig. 6b and Supplementary Table 10). Moreover, phylogenetic trees of C2H2 zinc finger and homeobox genes in the human genome revealed significantly expanded lineages containing a small number of previously annotated genes and a large number of newly annotated genes (Fig. 3c and Supplementary Fig. 6c). We also found remarkably expanded or newly constructed lineages of C2H2 zinc finger and homeobox gene families in other animal genomes including mouse (Supplementary Fig. 7).

### Validation and expression of the new gene model

We tested the accuracy of the new gene models by comparing the new gene models to UniProt^17^ and NR^18^ databases (Supplementary Fig. 8 and Supplementary Table 11-12). We determined that sequences in NR and UniProt databases matched a percentage of the newly annotated genes ranging from, 56% (C2H2 zinc finger) to 61% (NLR), and 39% (FAR1) to 57% (NLR), respectively, covering more than 80% of the sequences in those databases (Supplementary Fig. 8). For all gene families, the matched percentage of newly annotated genes was significantly lower than that of previously annotated genes because of relatively low coverage of *ab-initio* genes (Supplementary Fig. 8). However, we also found a large portion of previously annotated genes perfectly matched to the sequences in NR and UniProt, suggesting that many of the previously annotated genes were already present in those databases and these made higher coverage of the previously annotated genes. (Supplementary Fig. 9).

Swiss-Prot of UniProt is a database consisted of experimentally validated gene sequences^17^. In total, we detected that 25 and 1 newly annotated homeobox and C2H2 zinc finger genes in the human genome had identical sequences in Swiss-Prot, respectively (Supplementary Table 13). This may indicate that functional protein-coding genes were recorded in Swiss-Prot based on experimental evidences but yet annotated their gene structures from the genome sequences. When we compared previously annotated genes to SwissProt, the majority of perfectly matched genes to sequences of SwissProt was enriched in specific genomes such as human, mouse and Arabidopsis, suggesting that SwissProt primarily contained sequences of those species (Supplementary Fig. 10). Thus, we performed identification of new genes having strong homology to sequences in the SwissProt (>95% identity and 100% coverage) and found many newly annotated genes (65 and 34 of homeobox and C2H2 zinc finger, respectively) in animal genomes (Supplementary Fig. 11).

To validate the expression of the newly annotated genes, we performed RT-PCR and sequencing analyses using human, mouse and rice RNA samples (Fig. 4, Supplementary Fig. 12 and Supplementary Table 14). We designed 47, 43, and 79 primer sets for newly annotated genes in human, mouse and rice, respectively. Of these, 19, 15, and 50 genes (40, 35, and 63%) were confirmed expression and accurately validated by comparison between sequenced PCR products and annotated gene models. These results verify that the newly annotated genes are truly present and expressed as mRNA in each genome (Fig. 4 and Supplementary Table 14).

**Figure 4.**
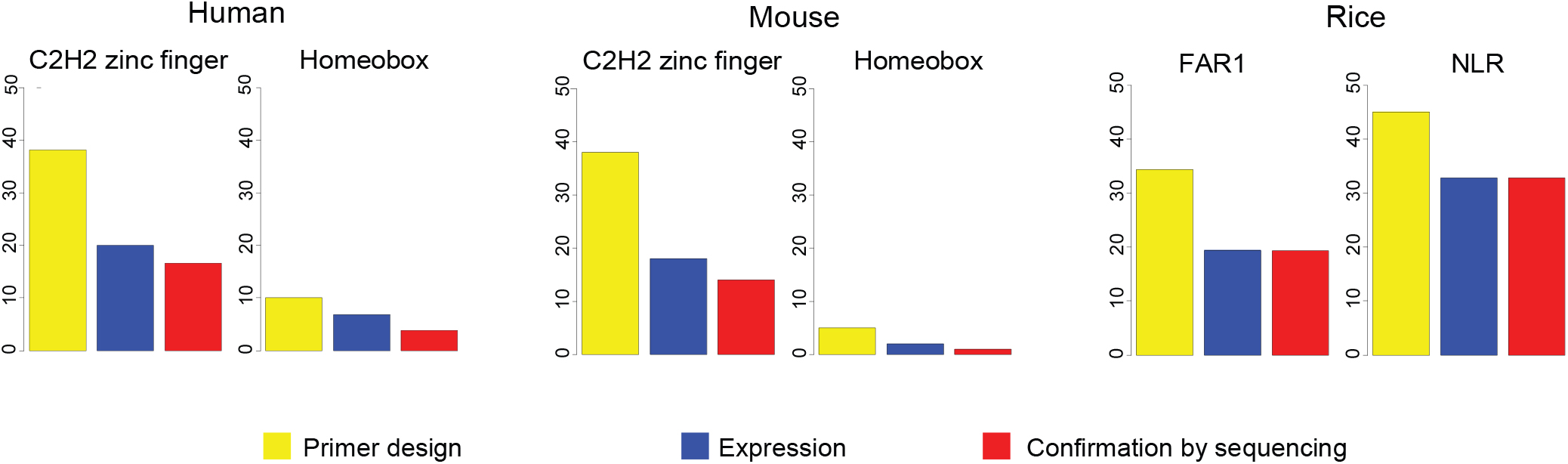
Validation and expression of newly annotated genes. Bar graphs show the numbers of primer-designed (yellow), expressed (blue) and validated genes (red) in human, mouse, and rice genomes, respectively, identified by RT-PCR and sequencing analyses.

### Annotation run-time using TGFam-Finder

The full annotation of a large genome can take weeks to complete and requires enormous computational resources^19^. To evaluate the performance of TGFam-Finder, we estimated the actual annotation run-time using TGFam-Finder for representative plant and animal genomes ranging from ~200 Mb to ~3 Gb using a desktop computer (32 Gb memory and 4 CPU). In total, it took 2 hours for NLRs of the *Selaginella* (~200 Mb) to 44 hours for C2H2 zinc finger of the human (~3 Gb) for annotation, indicating that completion of gene model construction could be finished within 2 days for the large genome such as human using a standard desktop computer (Table 1). Specifically, we verified that annotation of FAR1 in maize (~2 Gb) and C2H2 zinc finger in chicken (~1 Gb) required less than 12 hours. This indicates that users can efficiently annotate their target genes within half a day, with the exception of several huge genomes.

**Table 1.**
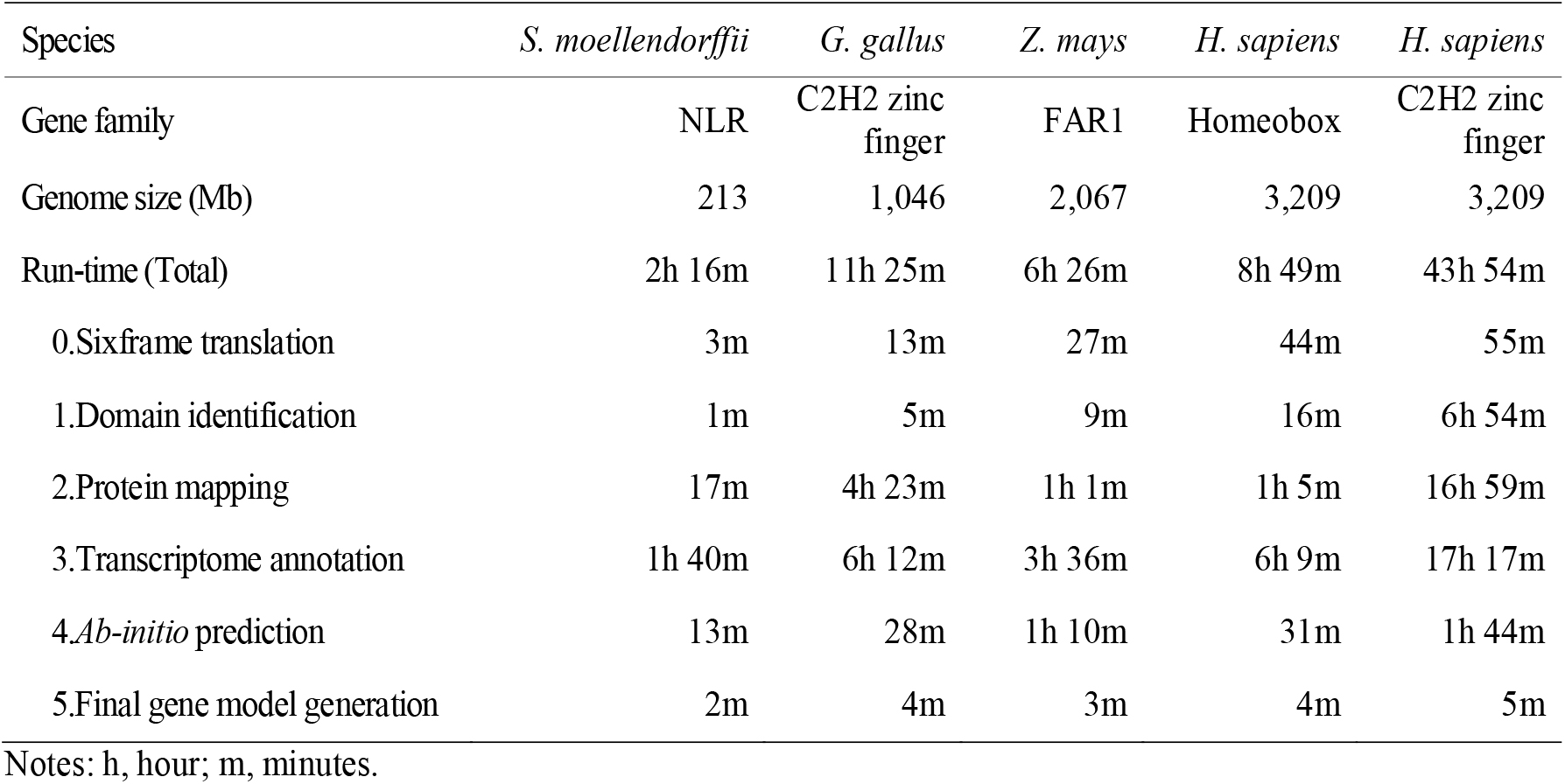
Annotation run-time of TGFam-Finder using a desktop PC.

| Species | *S. moellendorffii* | *G. gallus* | *Z. mays* | *H. sapiens* | *H. sapiens* |
| --- | --- | --- | --- | --- | --- |
| Gene family | NLR | C2H2 zinc finger | FAR1 | Homeobox | C2H2 zinc finger |
| Genome size (Mb) | 213 | 1,046 | 2,067 | 3,209 | 3,209 |
| Run-time (Total) | 2h 16m | 11h 25m | 6h 26m | 8h 49m | 43h 54m |
| 0.Sixframe translation | 3m | 13m | 27m | 44m | 55m |
| 1.Domain identification | 1m | 5m | 9m | 16m | 6h 54m |
| 2.Protein mapping | 17m | 4h 23m | 1h 1m | 1h 5m | 16h 59m |
| 3.Transcriptome annotation | 1h 40m | 6h 12m | 3h 36m | 6h 9m | 17h 17m |
| 4.*Ab-initio* prediction | 13m | 28m | 1h 10m | 31m | 1h 44m |
| 5.Final gene model generation | 2m | 4m | 3m | 4m | 5m |

Notes: h, hour; m, minutes.

### Genomic features of the newly annotated genes in plant and animal genomes

The genomic positions of the newly annotated genes could be classified into the following three categories: (1) non-overlapping, (2) overlapping with existing genes without target domain(s), and (3) overlapping with existing partial target genes (Supplementary Fig. 13). For non-overlapping genes, a large portion of those were overlapped with repetitive sequences (Supplementary Fig. 14a-b). Interestingly, the non-overlapping FAR1s in plant genomes were remarkably resided in the regions containing DNA-transposons. We observed that a significant number of NLRs in plant genomes co-localized with LTR-retrotransposons (Supplementary Fig. 14c). We also found that many C2H2 zinc finger were located in regions consisting of unclassified transposable elements in animal genomes. Considering previous reports describing annotation processes^19, 20^, our results suggest that repeat masking before gene annotation could have a crucial impact in generating imperfect gene models. In the case of newly annotated genes overlapping with existing genes without target domain(s), we observed that the newly annotated gene families were primarily overlapped with hypothetical genes without known domain(s). This result suggests that several newly annotated genes were ignored in previous annotations due to the presence of uncharacterized genes in the same region (Supplementary Fig. 15).

## Conclusion

The construction of accurate gene models after genome assembly is a critical step in genomic and functional analysis. Previous methods have been shown to be ineffective, hampered by imperfect methodologies, resources, and knowledge. Here, we described TGFam-Finder, a highly efficient tool to implement automatic structural annotation for target-gene families of interest. We evaluated and demonstrated the competitiveness of TGFam-Finder through the re-annotation of FAR1 and NLR gene families in 50 plants and C2H2 zinc finger and homeobox families in 50 animals. We only used publicly available resources, and identified large numbers of newly annotated genes that were omitted in the existing gene models (346, 45, 47, and 14% more FAR1, NLR, C2H2 zinc finger, and homeobox genes, respectively). The newly annotated genes were significantly supported by protein or transcriptome evidences. A total of 25 newly annotated homeobox and 1 C2H2 zinc finger genes in the human genome were identical to known functional gene sequences. We performed RT-PCR and sequencing analyses to confirm and validate the expression and accuracy of newly annotated genes in human, mouse, and rice genomes.

TGFam-Finder is easy to use and requires considerably less run-time and computing power than full annotation. The estimated annotation run-time of TGFam-Finder using a desktop PC was 2 hours for *Selaginella* (~200 Mb) NLRs and 44 hours for human (~3 Gb) C2H2 zinc finger. Compared to the long run-time of several weeks and intensive computational power required for full annotation, our results demonstrate that TGFam-Finder enables even novice users to obtain their target gene models within several days.

In summary, TGFam-Finder is designed to detect all protein-coding genes containing target domain(s) of interest and provide annotation evidence in eukaryotic genomes. TGFam-Finder enables users who study gene functions to determine their experimental priorities based on the annotation evidence and exact copy number of the genes of interest. Moreover, large-scale comparative studies of gene families will not be biased by missing genes, which are frequently encountered in previous annotations. Our approach provides an optimal solution for the identification and characterization of target-gene families, accelerating accurate functional, comparative and evolutionary analyses.

## Additional information

### Acknowledgements

This study was supported by the Basic Science Research Program through the National Research Foundation of Korea (NRF) funded by the Ministry of Education (NRF-2017R1A6A3A04004014) to S.K. and by a grant from the Agricultural Genome Center of the Next Generation Biogreen 21 Program of RDA (Project No. PJ013153) and by the National Research Foundation of Korea (NRF) grant funded by the Korea government(MSIT) (No. 2018R1A5A1023599, SRC) to D.C. We appreciate the assistance from the KOBIC Research Support Program. We also acknowledge to the following researchers who performed iterative beta testing of TGFam-Finder: Ho-Sub Shin, Myung-Shin Kim, and Jun-Ki Lee in Seoul National University; Namjin Koo in Korea Research Institute of Bioscience and Biotechnology; and Eunyoung Seo in University of California, Berkeley.

### Author contributions

S.K. and D.C. conceived the project, designed the content, and organized the manuscript. S.K., J.P., M.-S.K. and K.C. developed TGFam-Finder and annotation of the gene families. S.K., M.-S.K., Y.-M.K., and N.K. collected resources of 50 plant and 50 animal genomes. J.-H.K., S.-H.K., K.-S.K, N.O., S.-K.Y., and K.-S.P. prepared the RNA samples. J.-H.K., K.-S.K, N.O., and K.-S.P. implemented the RT-PCR analysis. S.K., M.-K.S., K.-T.K., J.J., H.K., Y.-Y.L., K.-H.S., H.C.M., and Y.-H.L performed phylogenetic analyses and validation of the new gene models. S.K. and H.K. designed and constructed the figures. S.K. and D.C. wrote the manuscript.

### URLs

GENECODE (human and mouse), https://www.gencodegenes.org/

TAIR (Arabidopsis), https://www.arabidopsis.org/

GenBank (Zebrafish), https://www.ncbi.nlm.nih.gov/genome/annotation_euk/Danio_rerio/105/

GenBank (Pig), https://www.ncbi.nlm.nih.gov/genome/annotation_euk/Sus_scrofa/106/

iTAG (Tomato), https://solgenomics.net/

RepeatModeler, http://www.repeatmasker.org/RepeatModeler/

RepeatMasker, http://www.repeatmasker.org/

## Methods

### Annotation overview of TGFam-Finder

TGFam-Finder was developed to run in the Linux OS environment. For novices in bioinformatics-based analyses, we constructed an install package that allow auto-installation of prerequisite tools to run TGFam-Finder (Supplementary Fig. 1). Through the install package, prerequisite tools, including Bowtie2-2.3.1^25^, HMMER-3.1b2^12^, BLAST 2.6.0+^26^, InterproScan-5.22-61.0^27^, Exonerate-2.2.0^21^, Blat v35^28^, Tophat-2.1.1 and Cufflinks-2.2.1^24^, Augustus-3.2.3^22^, Scipio-1.4^29^, and ClustalW-2.1^30^ are provided for further annotation using TGFam-Finder. To run TGFam-Finder, users need to configure the location information of genomic resources and the prerequisite programs in ‘RESOURCE.config’ and ‘PROGRAM_PATH.config’. Basically, ‘PROGRAM_PATH.config’ is automatically generated through the auto-installation process. Whereas, users should enter the location of the target genome, peptide sequences of target or allied species, and peptide sequences including target domains in various species as minimum resources. To classify specific proteins having target domain(s) of interest, TGFam-Finder requires the location of functional annotation information of target or allied species formatted as tsv and target domain ID(s) in ‘RESOURCE.config’. Moreover, users can input ‘EXTENSION_LENGTH’ to determine target regions for further annotation, and ‘MAX_INTRON_LENGTH’ for alignment processes using proteins. For extra configuration, users can also register the location of transcriptome, genomic position of genes and coding DNA sequences of target species in ‘RESOURCE.config’.

The annotation pipeline of TGFam-Finder consists of three steps. (1) Determination of target regions using ‘0.SixFrameTranslation.pl’ and ‘1.Domain_Identification.pl’, (2) Gene prediction in the target regions via ‘2.Auto_ProteinMapping.pl’, ‘3.Auto_ISGAP.pl’ and ‘4.Auto_Augustus.pl’, and (3) Generation of final gene model through ‘5. Generating_FinalGeneModel.pl’ (Fig. 1). To identify the position of target domains in an assembled genome, TGFam-Finder generates six-frame translated genome sequences. Then, a hidden Markov model (hmm) matrix is constructed through alignments among target domain(s) in protein sequences of target or allied species using ClustalW^30^. After identification of genomic regions containing target domain(s) using HMMER^12^, target regions including the target domain(s) and their flanking sequences are determined.

Structural annotation for the target regions is conducted via processes of protein mapping, transcriptome-based annotation and *ab initio* prediction. For efficient protein mapping, TGFam-Finder detects proteins with homology to target regions in the resource peptide sequences using BLAST+, and aligns between the proteins and matched target regions using Exonerate^21^. ‘Transcriptome-based annotation is implemented in the order of reference-guided transcriptome assembly using Tophat and Cufflinks^24^, and annotation via ISGAP pipeline^23^. For *ab-initio* gene prediction, the training set is constructed using the protein sequences of target or allied species having target domain(s), and the gene models generated from protein mapping and transcriptome-based annotation. Then, Augustus^22^ generates the gene model based on the training set. Finally, the final gene model is generated by combining the initial gene models in order from the transcriptome-based annotation, protein mapping, and *ab initio* prediction.

### Structural annotation of target-gene families in plant and animal genomes

FAR1 and NLR were re-annotated in 50 plant genomes, and C2H2 zinc finger and homeobox genes were re-annotated in 50 animal genomes using TGFam-Finder (Fig. 2-3, Supplementary Fig. 3 and 6, and Supplementary Table 3 and 7). We used assembled genomes and proteins described in Supplementary Table 1 and 2 as ‘TARGET_GENOME’ and ‘PROTEINS_FOR_DOMAIN_IDENTIFICATION’ in ‘RESOURCE.config’. After performing functional annotation using InterproScan-5^27^ for the proteins, generated tsv files for the proteins were used as ‘TSV_FOR_DOMAIN_IDENTIFICATION’. PF03101 (FAR1), PF00931 (NLR), PF00096 (C2H2 zinc finger), and PF00046 (homeobox) were selected as ‘TARGET_DOMAIN_ID’ for classification of target-gene families. ‘EXTENSION_LENGTH’ and ‘MAX_INTRON_LENGTH’ were determined as 30kb. We extracted target genes in the existing gene models of plants or animals having the Pfam IDs in the tsv files, and then merged them to use as ‘RESOURCE_PROTEIN’. Location of transcriptome, gff3, and coding DNA sequences of the plant and animal genomes were also recorded in ‘RESOURCE.config’ (Supplementary Table 1-2).

### Phylogenetic analyses of the new gene models

To perform phylogenetic comparison of new gene models for FAR1 and NLR in plant genomes, and C2H2 zinc finger and homeobox genes in animal genomes, we implemented gene clustering using OrthoMCL^31^ among the new models of each gene family (Supplementary Table 6 and 10). Then, the amino-acid sequences of target domains in each genome were aligned using ClustalW2^30^, and the phylogenetic trees of each gene family in the specific plant and animal genomes were constructed with MEGA7^32^ using the neighbor-joining method (Supplementary Fig. 4 and 7).

### Comparison of newly annotated genes and sequences in NR and UniProt

We compared new gene models of each gene family as query to sequences in NR and UniProt databases as subject using BLASTP. We counted the number of newly annotated genes that matched to NR and UniProt databases considering an e-value score of 1e^-5^ and more than 80% subject coverage (Supplementary Fig. 8 and Supplementary Table 11-12). To verify the number of newly annotated genes with strong homology to sequences in those databases, we used cut-off values greater than 98% identity and 100% subject coverage (Supplementary Fig. 9). We estimated the number of newly annotated genes with strong homology to functional genes in Swiss-Prot using cut-off values greater than 95% sequence similarity and 100% subject coverage (Supplementary Fig. 11).

### Validation of newly annotated genes using RT-PCR and sequencing analyses

MCF-7 and MCF-10A cell lines (ATCC, Teddington, UK) with LEC and ADSC (LONZA, Basel, Switzerland) were used for gene-expression verification and sequence validation for newly annotated genes in the human genome. Mouse embryonic fibroblasts (MEFs) (Stemgent, MA, USA) and two tissues (brain and spleen) from 6 to 8-week-old C57BL/6J mice (Koatech, Gyunggi-do, Korea) were used for analyses of the mouse genome. C57BL/6J mice were maintained according to the policies approved by CHA University. Seedlings of the rice (*Oryza sativa*) cultivar Nakdong were used for validation of rice FAR1 and NLR genes. Total RNA was extracted from rice seedlings, cultured human cells, and each mouse tissue using TRIzol^®^ reagent (Invitrogen, USA). The total RNA was treated with DNase I and reverse-transcribed with Oligo (dT) primers and Superscript II (Invitrogen), according to the manufacturer’s instructions. Subsequently, RT-PCR was conducted to analyze gene expression using the designed gene primers for each family. We used *GapdH* of human and mouse and *Actin* of rice as controls. PCR products were purified with AccuPrep^®^ PCR purification kit (Bioneer, Korea) or gel elution. Finally, the PCR products were sequenced using ABI3730XL (Applied Biosystems) as described in Supplementary Table 14.

### Identification of genes overlapping with repeat sequences

To annotate genomic sequences containing non-overlapping genes along with previously annotated genes described in Supplementary Figure 13, we performed repeat annotation for the genomic regions using RepeatModeler and RepeatMasker (see URL). *De novo* repeat libraries of each plant and animal genome were constructed using RepeatModeler, and then RepeatMasker was used to repeat masking on the repeat libraries. We considered that if specific repeat sequences covered more than 50% of a non-overlapping gene, the gene resided in the genomic region containing the specific repeat sequence (Supplementary Fig. 14).

### Computational resources used to run TGFam-Finder

To estimate annotation run-time of TGFam-Finder, we performed annotation using a desktop computer (Intel Core i7-4770 CPU @ 3.40GHz, 8 processors, and 32Gb memory) for NLR families in *S. moellendorffii*, C2H2 zinc finger gene families in *G. gallus*, FAR1 genes in *Z. mays*, and C2H2 zinc finger and homeobox genes in *H. sapiens* (Table 1). For efficient test, we randomly extracted and used ~10Gb of whole transcriptome data of *G. gallus*, *Z. mays*, and *H. sapiens as* described in Supplementary Table 1 and 2. For the annotation of FAR1 and NLR genes in 50 plant genomes, and C2H2 zinc finger and homeobox genes in 50 animal genomes, we used two of our computer servers with the following specifications (1) Intel Xeon CPU E5-2697 v2 @ 2.70GHz, 48 processors, and 264Gb memory, and (2) Intel Xeon CPU E5-4650 v2 @ 2.40GHz, 80 processors, and 512Gb memory. The re-annotation of each gene family was finished within one week using those servers.

## Data availability

The newly generated gene models including peptide and coding DNA sequences with gff3 and tsv are accessible at: http://TGFam-Finder.snu.ac.kr.

## Code availability

The TGFam-Finder program package including auto-installation and annotation scripts with sample data is accessible at: https://github.com/tgfam-finder and http://TGFam-Finder.snu.ac.kr.

